# Minimizing spectral overlap in multicolor flow cytometry experiments

**DOI:** 10.1101/2021.03.17.435861

**Authors:** Lars Rønn Olsen, Alfredo Benso, Gianfranco Politano, Mike Bogetofte Barnkob

## Abstract

Selecting fluorochromes for polychromatic panels for flow cytometry is complex and time-consuming. Poorly designed panels can result in overlap between the emission spectra of the different fluorochromes, making their signals difficult to separate. While assessing all possible panels is simple to do programmatically, the combinatorial complexity of most real world problems renders brute force computation impractical. Here we present a novel complexity optimization algorithm for fast design of minimal spectral overlap fluorochrome combinations. To aid researchers in designing fluorochrome panels we implemented the algorithm in a web server, Spectracular, that also considers instrument laser configuration and allows users to define various constraints related to antibody-fluorochrome availability and marker co-expression.

## Introduction

Fluorescent flow cytometry is a core methodology in cell biology research and clinical diagnostics. Primary usages include cell phenotyping, fluorescence-activated cell sorting, and sensitive validation of protein expression^1^. While these applications have traditionally been handled with panels of a dozen or fewer antibody-fluorochrome conjugates, advanced instruments with multiple lasers and detectors now enable the parallel measurement of up to 28-40 fluorochromes simultaneously^2,3^. This allows for novel biological insights into single cells in both health and disease^4^, but to truly leverage panels of this size, they must be designed with minimal overlap between the fluorochromes’ emission spectra.

Using multiple fluorescent proteins in the same panel will invariably increase the risk of emission spectra overlapping at unacceptable levels, and in turn lead to increased spill-over spreading error, which can confound the interpretation of the results^5^. Poor fluorochrome selections can also cause excessive background signal and errors when compensating post acquisition^6^, and ultimately lead to irreproducibility between instruments, users, and laboratories^7^. As flow cytometers incorporate more lasers and users design larger panels of antibodies, minimizing spectral spillover between fluorochromes becomes increasingly important but also increasingly complicated^8,9^.

While fluorochrome selection can be done manually by researchers when designing panels for a few antibodies, it quickly becomes time-consuming and technically challenging when including several antibodies^10^. In our experience, few flow cytometry users quantify the spectral overlap of their panels, but instead rely on online spectral viewers and personal experience. However, visually inspecting complex density curves is a poor way of identifying conflicting spectra, and humans perform suboptimally or even poorly, when attempting to solve combinatorial optimization problems of this sort^11^. The problem is NP hard as the combinatorial complexity increases approximately exponentially with size of the pool of fluorochromes to choose from. With new commercial fluorochromes being produced continuously, and the advent of spectral flow cytometry enabling even larger panels, the combinatorial complexity will continue to increase. To enable higher reproducibility and better panel design for flow cytometry, we have designed a complexity optimization algorithm to provide optimal or near-optimal fluorochrome combinations.

## Results

The optimal combination of spectra is here defined as the combination with the minimal mean overlap between fluorochromes. Selecting the optimal combination has a complexity of *O*(*n* choose *k*), where *n* is the number of fluorochromes to choose from and *k* is the number of desired fluorochromes in the panel. All solutions may be brute force evaluated when the number of operations, *N,* is small enough, thus providing the optimal solution. For cases with an impractically large *N*, the computational cost can be lowered by approximating the optimal solution. This is done by first clustering the spectra into *k* clusters, where each cluster contains highly overlapping spectra that constitute poor combinations in a panel. The clustered spectra can then be represented as a complete *k*-partite graph, by connecting vertices (representing fluorochromes) to all vertices not in the same cluster, where the edge weights correspond to the overlap coefficient. Then, the optimization problem can be formulated as a minimal maximum weight *k*-clique problem, which can be solved by removing all edges from the graph and iteratively adding them back starting from lowest to highest weight until a *k*-clique is formed. The approximation algorithm has a complexity of approximately *O*(*n*^3^/*k*) (**Supplementary Figure 1**). The approximation algorithm is outlined in **Figure 1**.

**Figure 1:**
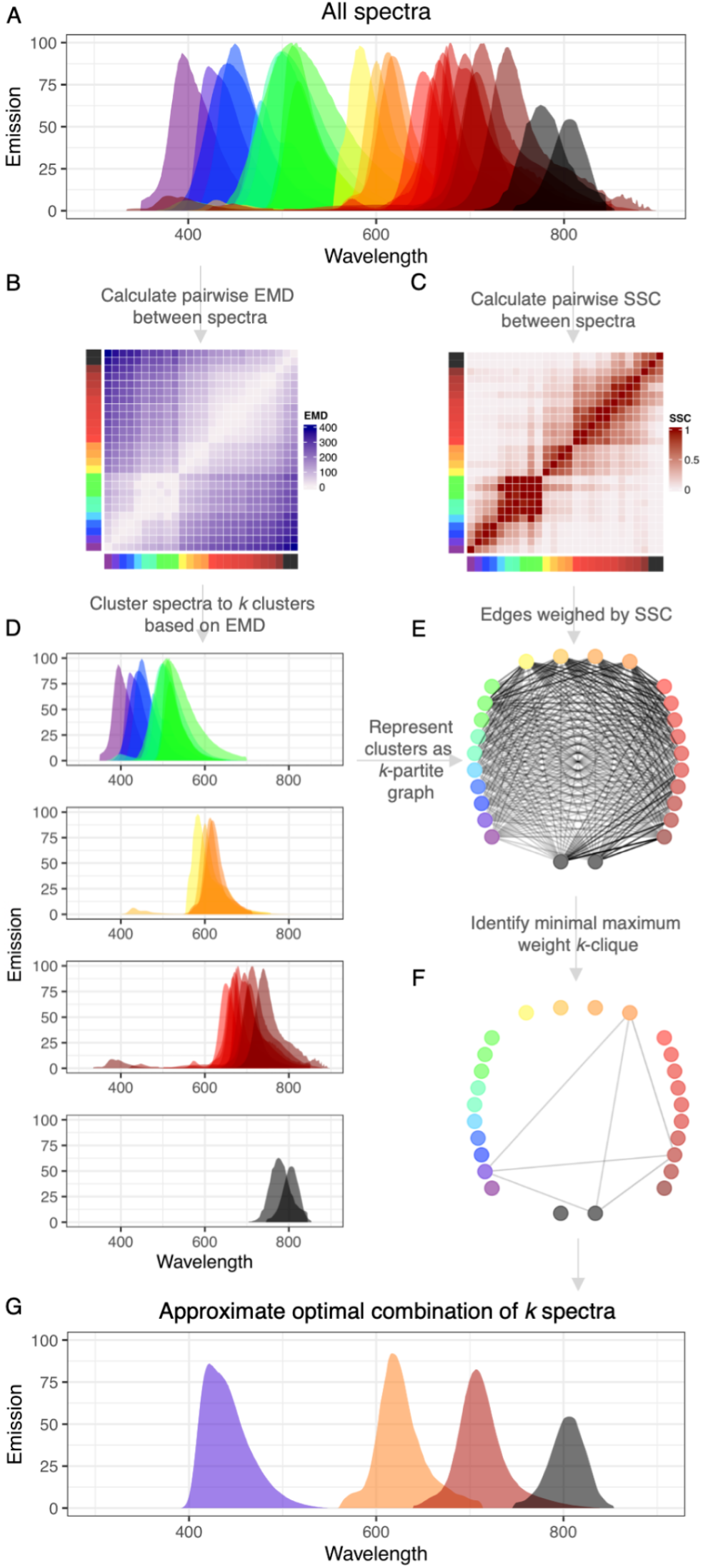
Outline of the complexity optimization algorithm. A) The emission spectra of all evaluated fluorochromes. B) Heatmap representing the pairwise earth mover’s distance between spectra. C) Heatmap representing the pairwise overlap coefficient between spectra. D) Emission spectra clustered to *k* clusters. E) The clusters represented as a *k*-partite graph, with edge weights corresponding to the overlap pairwise overlap coefficients. F) The minimal maximum weight *k*-clique, corresponding to G) the approximate optimal combination of *k* spectra.

In order to test the computational performance of the algorithm, we calculated the mean overlap coefficient for all spectra in 28 cases (30 choose [2;29]) and compared this to the brute force optimal solution, the brute force worst, and the mean of 100 random combinations. Spectracular provides optimal or near-optimal solutions to any *k* in the test (**Figure 2A**) and provides this solution in approximately 8 seconds (on a standard laptop, see materials and methods for details). In contrast, the brute force optimal solution quickly becomes computationally expensive to calculate as the combinatorial space increases, taking more than 41 minutes to calculate for *k* = 15 (**Figure 2B**). While this may be reasonable to some, in a case where a user would want to brute force compare 60 fluorochromes for a 28 multicolor panel, the computational power needed far exceeds what is available to most researchers (it would take approximately 50 years to calculate on a standard laptop). The approximate solution to this problem can be found in 2 minutes using Spectracular.

**Figure 2:**
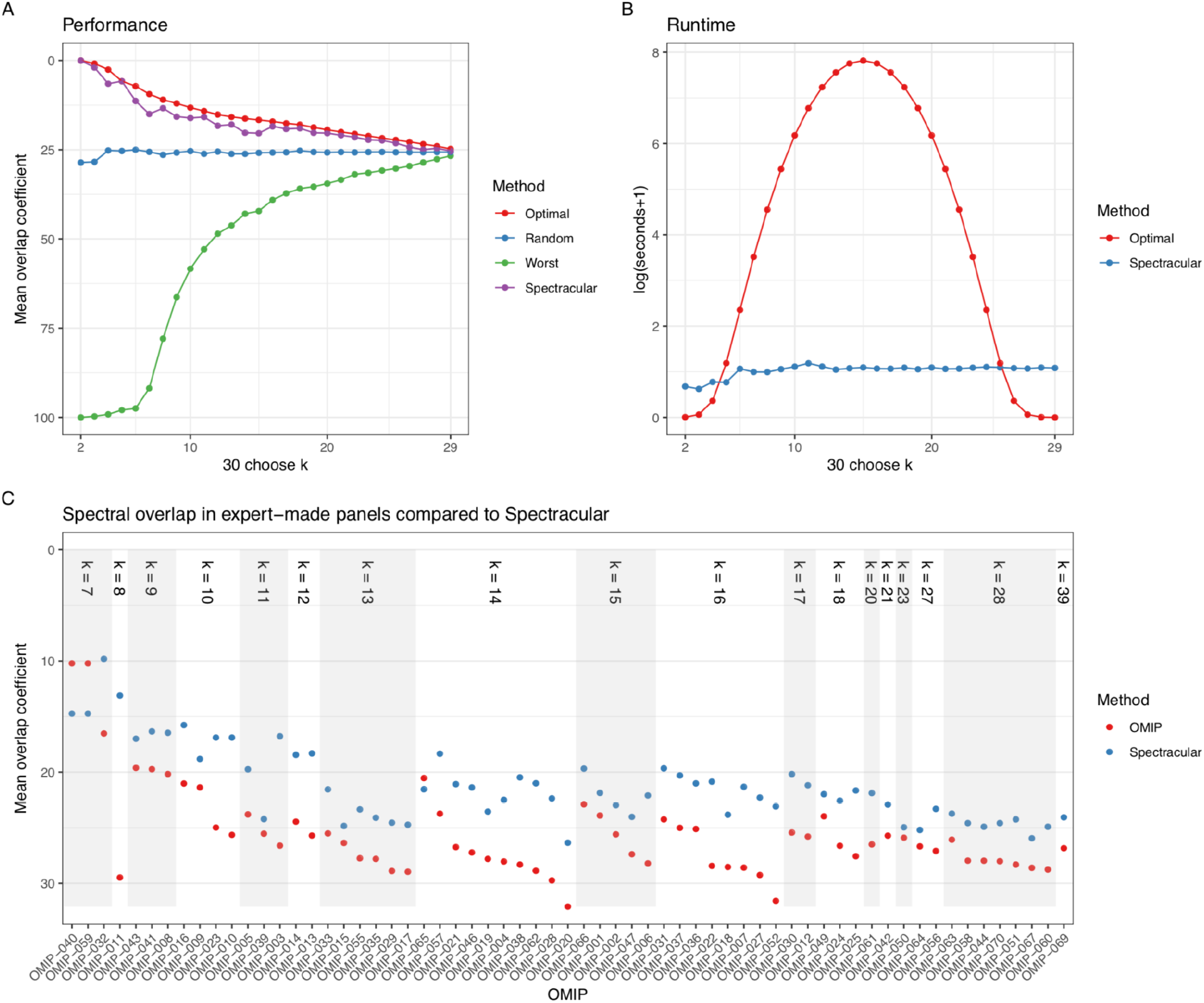
The performance and runtime (on a standard laptop, see materials and methods for details) of Spectracular on 30 choose 2-29 fluorochromes. A) Performance as measured by the mean overlap coefficient between all spectra in a given solution. The random method is the mean of 100 random combinations. B) Runtime of the brute force optimal solution and Spectracular in log(seconds+1). C) Analysis of the OMIP panels. The mean overlap coefficient of the fluorochromes in each OMIP was calculated and plotted in red. The mean overlap coefficient of the Spectracular solution of the same size was calculated and plotted in blue. OMIPs were ordered by the size of the panel.

Next, we wanted to compare panels suggested by our algorithm with panels designed and reviewed by flow cytometry experts. To this end, we analyzed 63 experiments described within the OMIP (Optimized Multicolor Immunofluorescence Panel) collection^3^. These panels are designed for instruments with 2-5 lasers to excite between 5-39 antibody-fluorochrome conjugates (see **Supplementary Table 1**). In the majority of cases we were able to identify a fluorochrome combination with a lower mean overlap coefficient (**Figure 2C**) in a matter of seconds. In a few cases the OMIP panels had lower mean overlap, which is likely due to the fact that our algorithm is designed to find the minimal maximum overlap combination, while the final solutions are evaluated by the mean overlap. The reason for this is that minimal maximum overlap panels inherently have a low mean overlap, while the opposite is not always true (**Supplementary Figure 2**). Still, comparing the mean overlap of panels is often more informative, as overlapping fluorochromes are sometimes inconsequential in real world applications - e.g. in the case of mutually exclusively expressed markers (see materials and methods for details).

Spectracular provides solutions to several flow cytometry panel design challenges. The basic functionality of the tool allows users to input the laser configuration of their flow cytometry instrument, how many antibodies they wish to detect, and which fluorochromes they want to choose from. Spectracular uses publicly available spectral data from multiple sources including the majority of fluorochromes and dyes commonly used in flow cytometry (currently 133 fluorochromes, but additional will be added as necessary). Based on this information, Spectracular approximates the combination of *k* fluorochromes, with minimal spectral overlap. Spectracular also offers more advanced functionalities, where users can pre-select fluorochromes (or groups of fluorochromes of which one is selected) to be included in the solution, after which an optimal panel revolving around the pre-selections is found (**Supplementary Figure 3**). This option can be useful when part of a panel has already been developed but a researcher needs to incorporate extra antibodies or wants to include an antibody that is only available with specific fluorochromes. It can also be used to leverage marker abundances and co-expressions, which can be highly useful to consider as fluorochromes conjugated to antibodies binding mutually exclusively expressed markers, are likely to never be excited on the same cells, and thus included in a panel with low risk of emission spillover.

Spectracular does not take into account users’ flow cytometer filter settings, brightness of fluorochromes, nor the natural decay of reagents such as tandem-dyes. These are all important factors to consider, but we specifically aimed for a simple, easy-to-use tool, which enables fast, near-optimal solutions to traditionally time consuming tasks. To our knowledge, Spectracular is the first tool that enables a quick, easy and near-optimal computational solution to the complex problem of designing fluorochrome panels. The tool is freely available for academic use at https://biosurf.org/spectracular.html.

## Materials and Methods

### Spectral data

Emission and excitation spectra were downloaded from Chroma Technology (www.chroma.com/), Semrock (www.semrock.com), and FPbase (www.fpbase.org/)^12^. In total, 133 commercially available fluorochromes were selected for the Spectracular web server. Based on the laser configuration submitted by the user, the emission spectra for all fluorochromes are calculated. A fluorochrome will be assumed to be excited by the laser which leads to the highest emission signal.

### Overlap and distance between spectra

The overlap between two emission spectra, *X* and *Y,* is calculated using the Szymkiewicz–Simpson coefficient (also known as the overlap coefficient), which is a similarity measure related to the Jaccard index. The overlap coefficient is calculated by dividing the integral of the intersection between two spectra, *X* and *Y*, with the integral of the smaller of the two spectra:

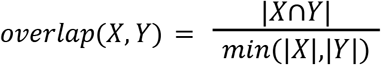

The distance between two spectra, *X*and *Y*, is calculated using the earth mover’s distance:

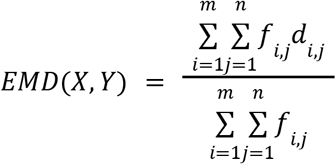

Where *m* and *n* are the number of bins of the two distributions (in Spectracular, emission spectra are binned to 100 bins), *f* is the flow between two bins, *i* and *j,* and *d* is the distance between two bins. The earth mover’s distances are then used to cluster spectra into *k* clusters using complete linkage hierarchical clustering.

### Performance evaluation

To evaluate the performance of Spectracular, we calculated the maximum overlap coefficient of spectra in a given solution, for *n* choose *k*, where *n*=30 and *k*=[2;29], using the following 30 spectra: BUV395, Brilliant Violet 421, Horizon V450, Pacific Blue, Brilliant Violet 480, Horizon V500, BUV496, BUV563, Brilliant Blue 515, Brilliant Violet 510, FITC, PE, Brilliant Violet 605, PE-Dazzle 594, PE-CF594, Brilliant Violet 650, APC, BUV661, PE-Cy5, PerCP-Cy5.5, Brilliant Blue 700, Brilliant Violet 711, BUV737, Brilliant Violet 750, PE-Cy7, PE-Fire 780, APC-H7, APC-Fire 750, Brilliant Violet 786, and BUV805 (we chose to perform this test on a subset of 30 fluorochromes, as increasing *n* will rapidly increase the combinatorial space to a size where it is simply not feasible to brute force calculate the optimal). For comparison with Spectracular, we calculated the brute force optimal solution, where the optimal solution for *k* fluorochromes was defined as the combination with the lowest maximum overlap coefficient. We also calculated the brute force worst solution, which was defined as the solution with the highest mean overlap coefficient. Finally we calculated the mean of 100 random solutions for each *k*. Similarly, we recorded the runtime for calculations of the brute force optimal solutions and the approximations when run on a 3.1 GHz Intel Core i7, 16 GB 2133 MHz LPDDR3 RAM, 2017 macbook pro, with computation parallelized on 7 cores.

### Comparison to expert-made panels

In order to compare results from Spectracular with human-made panels, we turned to the collection of Optimized Multicolor Immunofluorescence Panels (OMIPs) that have been published since 201013. This collection of flow cytometry panels are made and reviewed by flow cytometry experts and the methodology is described in great detail. We collected the fluorochrome panels and instrument laser configurations for 63 panels (see **Supplementary Table 1**) that were aimed at multicolor flow cytometry (excluding mass cytometry panels and panels without complete information on instrumentation, laser setup and fluorochrome usage). It should be noted that in our analysis we assume access to antibodies with any fluorochrome conjugate, which may not be the case in some real world applications.

### Optimization and evaluation metrics

Spectracular is designed to find the minimal maximum overlap fluorochrome combination, while the solutions are evaluated by the mean overlap. The reason for optimizing by minimizing the maximum overlap coefficient is that a low mean overlap coefficient alone does not guarantee a useful panel if, for example, a pair of fluorochromes in the solution has an unacceptably large overlap. However, minimizing the maximum overlap coefficient guarantees that no two fluorochromes overlap to an unacceptable degree, while invariably resulting in a low mean overlap coefficient (**Supplementary Figure 2**). The reason for evaluating solutions by their mean overlap coefficient, is that many real world panels are purposely designed with pairs of overlapping fluorochromes. This is a useful strategy if the fluorochromes are conjugated to antibodies targeting proteins that are mutually exclusively expressed on single cells, and thus the fluorochromes will never be excited simultaneously. In these cases, the minimal maximum overlap coefficient is a poor indicator of the quality of the panel, while mean overlap coefficient provides a better, albeit not perfect metric of panel quality.

### Implementation

Spectracular was implemented in R14, and the graphical user interface was developed using Shiny^15^. While users can currently choose from 133 different fluorochromes, a list of 55 pre-selected fluorochromes are suggested as defaults. These were selected by comparing the emission overlap between all fluorochromes and choosing the ones with least overall emission overlap. A minimal excitation ratio for proposed fluorochromes is set to 25%, but can be changed by the user. This will also impact the calculation of the overlap coefficient, as overlaps below the minimal excitation ratio are ignored.

## Code availability

The standalone R script to run Spectracular locally can be downloaded at https://github.com/biosurf/spectracular.

## Acknowledgements

We would like to thank Peter Wad Sackett, Leon Eyrich Jessen, and Kristoffer Vitting-Seerup for valuable discussions about the complexity optimization algorithm, and Christina Bligaard Pedersen for valuable input to the visual representations.

## Author contributions

LRO and MBB conceived of the study. LRO designed and implemented the complexity optimization algorithm with input from AB and GP. LRO, AB, and GP designed and implemented the advanced inclusion options. LRO implemented the Shiny server. LRO and MBB conceived of, and carried out the performance evaluation study. LRO and MBB drafted the manuscript. All authors critically reviewed the manuscript and approved of the submitted version.

