## Supplementary material for "Minimizing spectral overlap in multicolor flow cytometry experiments": supplementary materials.pdf

### Supplementary figures

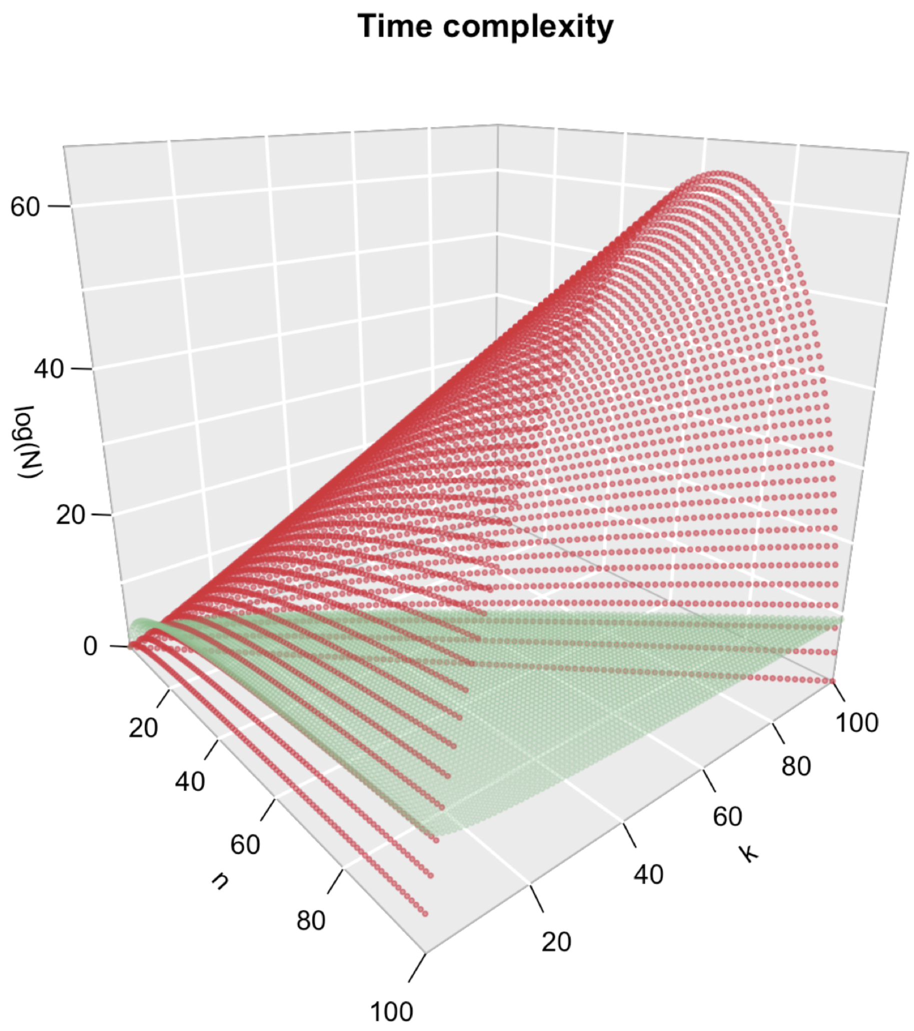


**Supplementary Figure 1.** Time complexity of algorithms for selecting minimal overlap panels. Red: "brute force" evaluation of all possible combinations at O(*n* choose *k*). Green: approximation using Spectracular at O($n^3/k$). Axes are *n*: number of fluorochromes to choose from, *k*: number of fluorochromes in a given solution, *log(N)*: logarithm of the operations required to solve the problem, *N*.

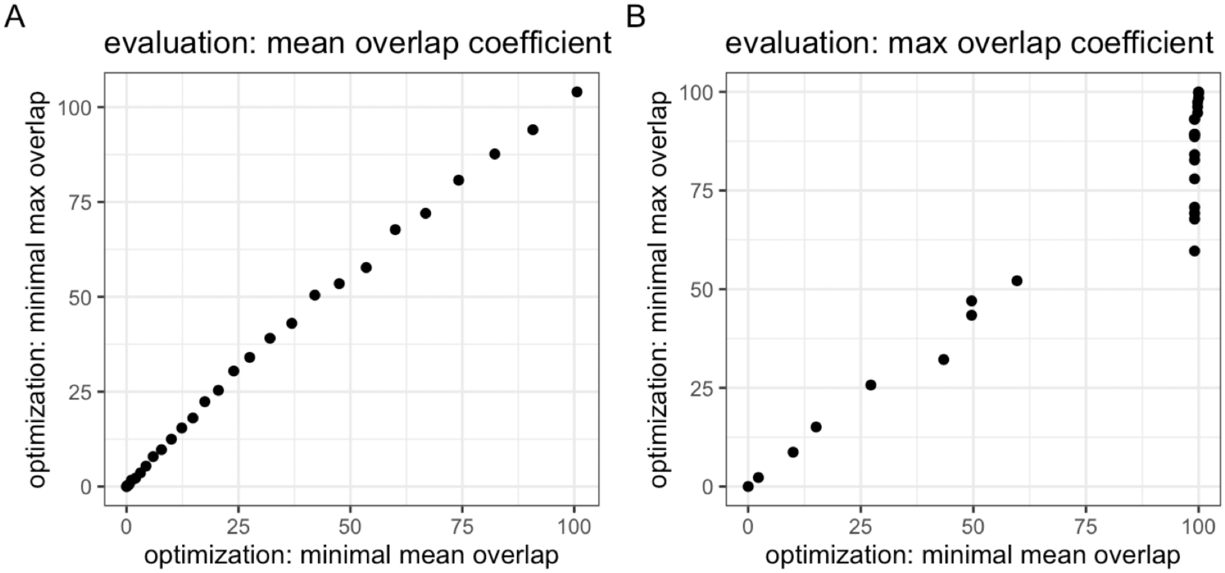


**Supplementary Figure 2.** Comparison of two different optimization and evaluation metrics. A) mean overlap coefficient of 100 choose [1;100] optimized by minimizing mean overlap coefficient (x axis) and max overlap between pairs of fluorochromes (y axis). As seen, the mean overlap coefficient of the solutions varies only slightly between optimizations methods. B) Same as A, except solutions are evaluated by the maximum overlap coefficient. As can be seen, solutions optimized by minimizing the mean overlap coefficient of the solutions (x axis), will often include pairs of fluorochromes for which the emission spectra overlap by almost 100%. When optimizing to minimize the maximum pairwise overlap coefficient (y axis), this is of course not the case.

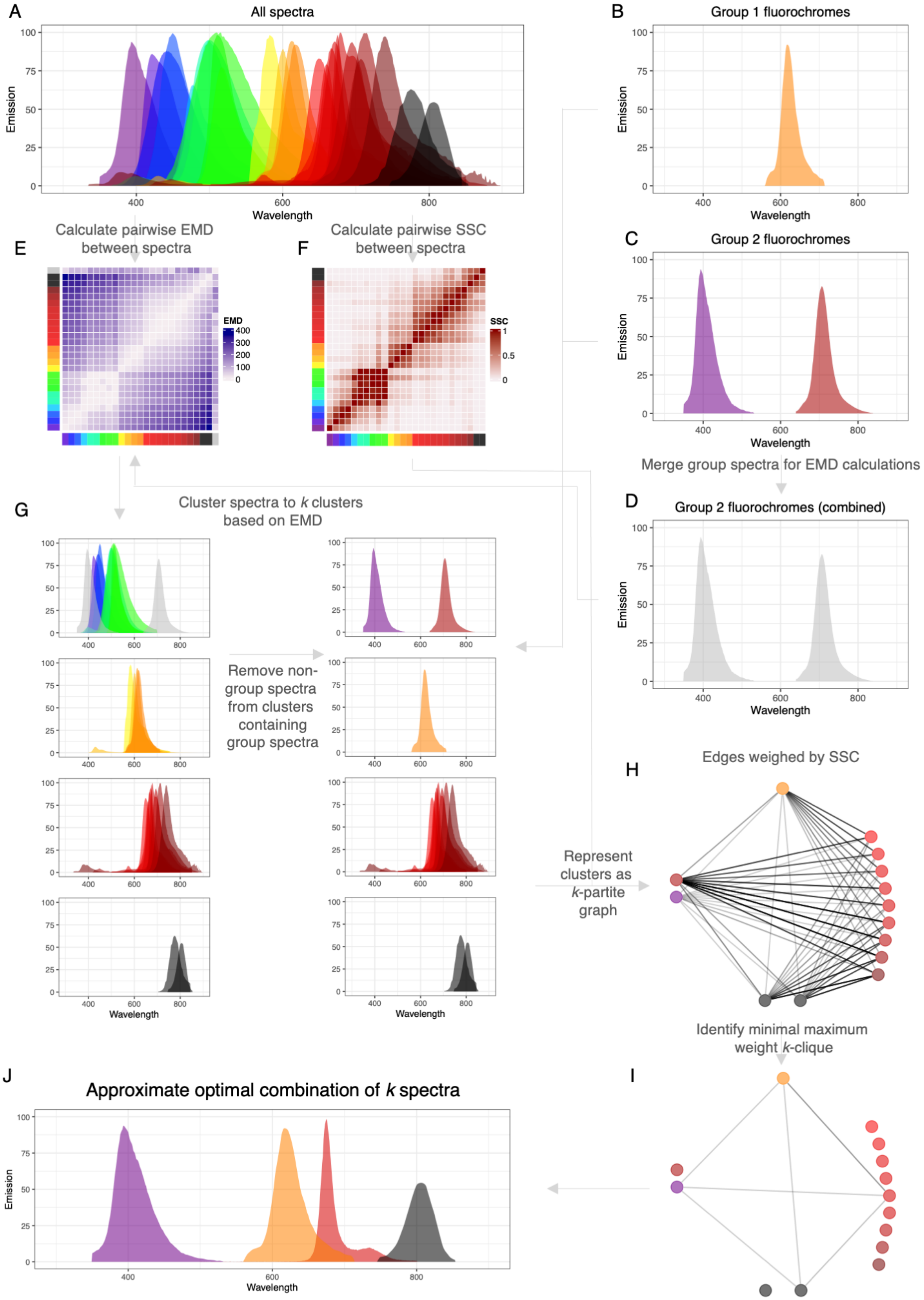


Legend on next page.

**Supplementary Figure 3.** Overview of run 26 choose 4 with two pre-selected groups of fluorochromes. A) All spectra to design panel from. B) Group 1 pre-selection which consists of one fluorochrome that must be included in the final solution. C) Group 2 pre-selection which consists of two fluorochromes of which precisely one must be included in the final solution. D) Group 2 fluorochromes merged into one spectrum for the purpose of calculating the pairwise earth mover's distance matrix. E) Matrix with pairwise earth mover's distances for all fluorochromes or groups. F) Matrix with pairwise overlap coefficients for all fluorochromes. G) Clustered spectra and groups based on earth mover's distance (left), and the clusters with groups expanded to individual spectra as well as non-group spectra removed from the clusters containing group spectra. H) *k*-partite graph where partites corresponds to clusters, nodes corresponds to spectra, and edges are weighted by the overlap coefficients. I) Minimal maximum weight *k*-clique corresponding to J) the approximate optimal combination of 4 spectra.

### Supplementary tables

Supplementary Table 1. OMIP panel name, laser setup and fluorochrome usage which Spectracular was tested against.

| OMIP | Lasers (nm) | Fluorochromes used |
|---|---|---|
| OMIP-001 | 633, 534, 488, 407 | APC, PE-Cy7, PE-Cy5, PE, Alexa Fluor 594, Alexa Fluor 488, LIVE-DEAD Fixable Violet, Pacific Blue, APC-Cy7, Alexa Fluor 660, Qdot 705, Qdot 655, Qdot 605, Qdot 585, Qdot 545 |
| OMIP-002 | 633, 534, 488, 407 | APC-R700, APC, PE-Cy5, PE, FITC, Pacific Blue, LIVE-DEAD Fixable Violet, APC-Cy7, PE-Alexa Fluor 750, PE-Alexa Fluor 700, PE-Alexa Fluor 594, Qdot 800, Qdot 705, Qdot 605, Qdot 565 |
| OMIP-003 | 488, 633, 405 | PE-Cy7, PerCP-Cy5.5, PE, FITC, APC, Pacific Blue, PE-Alexa Fluor 610, APC-Cy7, Alexa Fluor 680, Qdot 605, Pacific Orange |
| OMIP-004 | 633, 532, 488, 407 | Horizon V450, PE-Cy7, PerCP-Cy5.5, PE-Texas Red, PE-Cy5, Alexa Fluor, LIVE-DEAD Fixable Aqua 647, Alexa Fluor 700, PE-Cy5.5, PE, Qdot 800, APC-Cy7, Qdot 605, Qdot 655 |
| OMIP-005 | 633, 534, 488, 408 | APC, PE-Cy7, PE-Cy5, PE, FITC, Pacific Blue, LIVE-DEAD Fixable Aqua, PE-Texas Red, Alexa Fluor 680, APC-Cy7, Qdot 605 |
| OMIP-006 | 635, 532, 488, 407 | Alexa Fluor 700, Alexa Fluor 647, PE-Cy7, PE-Cy5.5, PE-Cy5, PE-Texas Red PerCP-Cy5.5, FITC, Horizon V500, APC-Cy7, Qdot 705, Qdot 655, Qdot 605, Qdot 585, Qdot 565, Qdot 545 |
| OMIP-007 | 488, 405, 633, 532 | PerCP-Cy5.5, FITC, Pacific Blue, APC-H7, APC, Alexa Fluor 700, LIVE-DEAD Fixable Aqua, PE-Cy7, PE-Cy5.5, PE-Cy5, PE-Texas Red, PE, Qdot 800, Qdot 705, Qdot 655, Qdot 605 |
| OMIP-008 | 633, 488, 405 | APC, PE-Cy7, PerCP-Cy5.5, PE, Pacific Blue, Alexa Fluor 488, Alexa Fluor 700, Qdot 605, APC-Cy7 |
| OMIP-009 | 633, 534, 488, 408 | APC, PE-Cy7, PE-Cy5, PE, FITC, Pacific Blue, LIVE-DEAD Fixable Aqua, PE-Texas Red, Alexa Fluor 680, APC-Cy7 |

| | | |
|---|---|---|
| OMIP-010 | 488, 638, 405 | FITC, PE, PE-Texas Red, PE-Cy5.5, PE-Cy7, APC, APC-Alexa Fluor 700, APC-Alexa Fluor 750, Pacific Blue, Krome Orange |
| OMIP-011 | 633, 488 | LIVE-DEAD Fixable Near-IR, Alexa Fluor 700, Alexa Fluor 647, PE-Cy7, PE-Cy5, PE-Texas Red, PE, SYTO 16 |
| OMIP-012 | 640, 532, 488, 406 | APC-Cy7, Alexa Fluor 700, APC, PE-Cy7, PE-Cy5.5, PE-Cy5, PE-Texas Red, PE, PerCP, FITC, Qdot 705, Qdot 655, Qdot 605, Qdot 585, Qdot 565, AmCyan, Pacific Blue |
| OMIP-013 | 628, 532, 488, 405 | APC-H7, Alexa Fluor 680, APC, PE-Cy7, PE-Cy5.5, PE-Cy5, PE, FITC, Qdot 705, Qdot 605, Brilliant Violet 421, LIVE-DEAD Fixable Aqua |
| OMIP-014 | 633, 532, 488, 405 | APC-Cy7, Alexa Fluor 700, APC, PE-Cy7, PE-Cy5, PE-Texas Red, PE, PerCP-Cy5.5, FITC, Qdot 655, LIVE-DEAD Fixable Aqua, Pacific Blue |
| OMIP-015 | 628, 532, 488, 405 | APC-eFluor 780, Alexa Fluor 680, PE-Cy7, PE-Cy5, PE-CF594, PE, FITC, Brilliant Violet 786, Qdot 655, Qdot 605, Qdot 585, LIVE-DEAD Fixable Aqua, Brilliant Violet 421 |
| OMIP-016 | 633, 488, 408, 355 | APC-Cy7, Alexa Fluor 700, APC, PE-Cy7, PerCP-Cy5.5, PE, FITC, Horizon V500, Horizon V450, LIVE-DEAD Fixable Blue |
| OMIP-017 | 628, 532, 488, 405 | Alexa Fluor 680, Alexa Fluor 647, PE-Cy7, PE-Cy5, Alexa Fluor 594, PE, FITC, Qdot 800, Qdot 655, Brilliant Violet 605, Qdot 585, LIVE-DEAD Fixable Aqua, Brilliant Violet 421 |
| OMIP-018 | 633, 561, 488, 405 | APC-Cy7, Alexa Fluor 700, APC, PE-Cy7, PE-Cy5.5, PE-Cy5, PE-Texas Red, PE, PerCP, FITC, Qdot 800, Qdot 700, Qdot 655, Qdot 605, Brilliant Violet 510, Pacific Blue |
| OMIP-019 | 628, 532, 488, 405 | APC-Cy7, Alexa Fluor 680, APC, PE-Cy5.5, PE-Cy5, Alexa Fluor 594, PE, FITC, Brilliant Violet 786, Qdot 655, Brilliant Violet 605, Qdot 585, LIVE-DEAD Fixable Aqua, Brilliant Violet 421 |
| OMIP-020 | 405, 488, 633 | Brilliant Violet 711, Brilliant Violet 650, Brilliant Violet 605, Pacific Orange, Pacific Blue, PE-Cy7, PerCP-Cy5.5, Per-CP, PE-Texas Red, PE, FITC, APC-H7, Alexa Fluor 700, APC |

| | | |
|---|---|---|
| OMIP-021 | 405, 488, 561, 633 | Brilliant Violet 421, LIVE-DEAD Fixable Aqua, Brilliant Violet 605, Brilliant Violet 650, Brilliant Violet 711, Brilliant Violet 786, FITC, PerCP-Cy5.5, PE, PE-Cy5, PE-Cy7, APC, Alexa Fluor 700, APC-Cy7 |
| OMIP-022 | 488, 407, 640, 532 | PerCP-Cy5.5, Alexa Fluor 488, Brilliant Violet 786, Brilliant Violet 711, Brilliant Violet 650, Brilliant Violet 570, Qdot 565, LIVE-DEAD Fixable Aqua, Horizon V450, APC-eFluor 780, Alexa Fluor 700, APC, PE-Cy7, PE-Cy5.5, PE-Texas Red, PE |
| OMIP-023 | 488, 638, 405 | FITC, PE, PE-Texas Red, PE-Cy5.5, PE-Cy7, APC, APC-Alexa Fluor 700, APC-H7, Pacific Blue, Horizon V500 |
| OMIP-024 | 628, 532, 488, 405, 355 | APC-Cy7, Alexa Fluor 700, APC, PE-Cy7, PE-Cy5, PE-Texas Red, PE, PerCP-Cy5.5, FITC, Brilliant Violet 786, Brilliant Violet 711, Brilliant Violet 650, Brilliant Violet 605, Brilliant Violet 570, Brilliant Violet 421, LIVE-DEAD Fixable Aqua, BUV737, BUV395 |
| OMIP-025 | 628, 532, 488, 405, 355 | APC-H7, Alexa Fluor 700, APC, PE-Cy7, PE-Cy5, PE-eFluor 610, PE, PerCP-Cy5.5, FITC, Brilliant Violet 786, Brilliant Violet 711, Brilliant Violet 650, Brilliant Violet 605, Brilliant Violet 570, Brilliant Violet 421, LIVE-DEAD Fixable Aqua, BUV737, BUV395 |
| OMIP-027 | 488, 405, 633, 532 | PerCP-eFluor 710, FITC, Brilliant Violet 786, Brilliant Violet 711, Brilliant Violet 650, Brilliant Violet 605, LIVE-DEAD Fixable Aqua, Pacific Blue, APC-H7, Alexa Fluor 700, APC, PE-Cy7, PE-Cy5.5, PE-Cy5, PE-Texas Red, PE |
| OMIP-028 | 405, 488, 633 | Brilliant Violet 786, Brilliant Violet 711, Brilliant Violet 650, Brilliant Violet 605, Brilliant Violet 510, Brilliant Violet 421, PE-Cy7, PE-Cy5, PE-Texas Red, PE, FITC, LIVE-DEAD Fixable Near-IR, Alexa Fluor 700, Alexa Fluor 647 |
| OMIP-029 | 628, 532, 488, 405 | APC-H7, Alexa Fluor 680, Alexa Fluor 647, PE-Cy7, PE-Cy5.5, PE-Cy5, PE-CF594, PE, FITC, Brilliant Violet 605, Brilliant Violet 570, Brilliant Violet 421, LIVE-DEAD Fixable Aqua |
| OMIP-030 | 355, 405, 488, 561, 640 | LIVE-DEAD Fixable Blue, eFluor 450, Brilliant Violet 510, Brilliant Violet 570, Brilliant Violet 711, Brilliant Violet 786, Qdot 605, eFluor 660, Alexa Fluor 488, PerCP-Cy5.5, PE, PE-CF594, PE-Cy5.5, PE-Cy7, APC, Alexa Fluor 700, APC-eFluor 780 |

| | | |
|---|---|---|
| OMIP-031 | 488, 405, 640, 561, 355 | PerCP-Cy5.5, Alexa Fluor 488, Brilliant Violet 786, Brilliant Violet 711, Brilliant Violet 605, Brilliant Violet 510, DAPI, APC-Cy7, Alexa Fluor 700, APC, PE-Cy7, PE-CF594, PE, BUV805, BUV737, BUV395 |
| OMIP-032 | 405, 488, 561, 640 | APC, APC-Cy7, Brilliant Violet 421, FITC, PE, PE-Cy7, PerCP-Cy5.5 |
| OMIP-033 | 640, 532, 488, 405 | APC-Cy7, Alexa Fluor 700, APC, PE-Cy7, PE-Texas Red, PE, PerCP-Cy5.5, FITC, Brilliant Violet 786, Brilliant Violet 711, Brilliant Violet 605, Brilliant Violet 510, Brilliant Violet 421 |
| OMIP-035 | 405, 488, 633 | LIVE-DEAD Fixable Near-IR, Brilliant Violet 786, PE-CF594, Brilliant Violet 711, Pacific Blue, PerCP-Cy5.5, PE-Cy7, PE, APC, FITC, Alexa Fluor 700, PE, Brilliant Violet 510 |
| OMIP-036 | 488, 405, 640, 532, 355 | LIVE-DEAD Fixable Aqua, Brilliant Violet 510, BUV395, BUV496, BUV805, Brilliant Violet 786, Brilliant Blue 515, APC-H7, Brilliant Violet 650, Alexa Fluor 647, Brilliant Violet 711, Brilliant Violet 605, PE-Cy7, PerCP-Cy5.5, Brilliant Violet 421, PE-CF594 |
| OMIP-037 | 641, 488, 402, 355 | BUV395, BUV737, BUV805, Brilliant Violet 421, Brilliant Violet 510, Brilliant Violet 605, Brilliant Violet 650, Brilliant Violet 786, Alexa Fluor 488, PE, PE-eFluor 610, PE-eFluor 710, PE-Cy7, APC, Alexa Fluor 700, APC-Cy7 |
| OMIP-038 | 640, 532, 488, 405 | eFluor 780, Alexa Fluor 700, APC, PE-Cy7, PE-CF594, PE, PerCP-eFluor 710, FITC, Brilliant Violet 786, Brilliant Violet 711, Brilliant Violet 650, eFluor 605, Horizon V500, eFluor 450 |
| OMIP-039 | 628, 561, 488, 405 | APC-Vio770, APC, PE-Vio770, PE-Cy5, PE-Dazzle 594, PE, PerCP-Cy5.5, Brilliant Violet 786, Brilliant Violet 605, LIVE-DEAD Fixable Aqua, eFluor 450 |
| OMIP-040 | 488, 561, 640, 405, 355 | BUV395, Brilliant Violet 421, APC, APC-Cy7, PE, Brilliant Blue 515, PerCP-Cy5.5 |
| OMIP-041 | 405, 488, 633 | APC-Cy7, PE-Cy7, Brilliant Violet 421, PE, APC, PE, Alexa Fluor 647, PerCP-Cy5.5, Ghost Dye Violet 510 |
| OMIP-042 | 355, 405, 488, 561, 640 | BUV395, LIVE-DEAD Fixable Blue, BUV496, BUV661, BUV737, Brilliant Violet 421, Brilliant Violet 450, Brilliant Violet 510, Brilliant Violet 605, Brilliant Violet 650, Brilliant Violet 711, Brilliant Violet 786, Alexa Fluor 488, PerCP-Cy5.5, PE, |

| | | |
|---|---|---|
| | | PE-CF594, PE-Cy5, PE-Cy7, APC, Alexa Fluor 700, APC-eFluor 780 |
| OMIP-043 | 488, 405, 355, 640, 532 | Brilliant Violet 421, Brilliant Violet 786, APC, BUV395, PerCP-Cy5.5, Brilliant Violet 510, FITC, PE, LIVE-DEAD Fixable Blue |
| OMIP-044 | 355, 405, 488, 532, 628 | BUV395, LIVE-DEAD Fixable Blue, BUV496, BUV563, BUV661, BUV737, BUV805, Brilliant Violet 421, Brilliant Violet 480, Brilliant Violet 570, Brilliant Violet 605, Brilliant Violet 650, Brilliant Violet 711, Brilliant Violet 750, Brilliant Violet 786, FITC, Brilliant Blue 630, Brilliant Blue 660, Brilliant Blue 700, Brilliant Blue 790, PE, PE-CF594, PE-Cy5, PE-Cy5.5, PE-Cy7, Alexa Fluor 647, Alexa Fluor 700, APC-H7 |
| OMIP-046 | 488, 406, 640, 532 | FITC, PerCP-Cy5.5, Brilliant Violet 786, Brilliant Violet 711, Brilliant Violet 650, Brilliant Violet 605, Pacific Blue, LIVE-DEAD Fixable Aqua, APC-H7, Alexa Fluor 700, APC, PE-Cy7, PE-Texas Red, PE |
| OMIP-047 | 405, 488, 561, 640 | APC-Cy7, LIVE-DEAD Fixable Near-IR, Brilliant Violet 786, Brilliant Violet 650, Alexa Fluor 700, Brilliant Violet 711, PE-CF594, PE, FITC, APC, Brilliant Violet 605, PE-Cy7, Brilliant Violet 510, Brilliant Violet 421, PerCP-eFluor 710 |
| OMIP-049 | 488, 405, 640, 561, 355 | PerCP-Cy5.5, Brilliant Blue 515, Brilliant Violet 786, Brilliant Violet 711, Brilliant Violet 650, Brilliant Violet 605, Brilliant Violet 510, Brilliant Violet 421, APC-Cy7, Alexa Fluor 700, APC, PE-Cy7, PE-CF594, PE, BUV805, BUV737, BUV496, BUV395 |
| OMIP-050 | 488, 408, 532, 355, 628 | Brilliant Blue 515, Brilliant Blue 630, Brilliant Blue 660, Brilliant Blue 700, Brilliant Blue 790, Brilliant Violet 421, Brilliant Violet 480, Brilliant Violet 570, Brilliant Violet 605, Brilliant Violet 650, Brilliant Violet 711, Brilliant Violet 750, Brilliant Violet 786, BUV395, BUV496, BUV563, BUV661, BUV737, BUV805, APC, APC-R700, APC-Fire 750, LIVE-DEAD Fixable Blue |
| OMIP-051 | 355, 405, 488, 532, 628 | Brilliant Blue 515, Brilliant Blue 630, Brilliant Blue 660, Brilliant Blue 700, Brilliant Blue 790, PE, PE-Dazzle 594, PE-Cy5, PE-Cy5.5, PE-Cy7, APC, APC-R700, APC-H7, BUV395, BUV496, BUV563, BUV661, BUV737, BUV805, LIVE-DEAD Fixable Blue, Brilliant Violet 421, Brilliant Violet 510, Brilliant Violet 570, Brilliant Violet 605, Brilliant Violet 650, Brilliant Violet 711, Brilliant Violet 750, Brilliant Violet 786, |

| | | |
|---|---|---|
| OMIP-052 | 628, 532, 488, 408 | LIVE-DEAD Fixable Aqua, APC-Cy7, PE-Cy5.5, Brilliant Violet 570, PE-Texas Red, Alexa Fluor 700, Brilliant Violet 750, Brilliant Violet 421, Brilliant Violet 605, Alexa Fluor 647, PE-Cy5, Brilliant Violet 650, Brilliant Violet 711, PE, Brilliant Violet 785, PE-Cy7 |
| OMIP-055 | 638, 488, 405 | APC-Cy7, APC, Alexa Fluor 700, PE-Dazzle 594, PE, FITC, PE-Cy5, PE-Cy7, Brilliant Violet 421, Brilliant Violet 510, Brilliant Violet 605, Brilliant Violet 650, Brilliant Violet 786 |
| OMIP-056 | 637, 532, 488, 405, 355 | BUV805, Horizon V450, LIVE-DEAD Fixable Blue, Brilliant Blue 660, BUV395, PE-Cy7, PE-Cy5.5, BUV563, Brilliant Violet 786, Brilliant Violet 750, APC, Brilliant Blue 700, FITC, PE, BUV496, PE-Cy5, Brilliant Violet 605, PE-Vio615, APC, Alexa Fluor 700, APC-Vio770, Brilliant Violet 711, BUV661, BUV737, Brilliant Violet 480, Brilliant Violet 570, Brilliant Violet 650 |
| OMIP-057 | 355, 405, 488, 561, 640 | PI (Propidium iodide), BUV395, BUV737, Brilliant Violet 421, Brilliant Violet 510, Brilliant Violet 605, Brilliant Violet 786, FITC, PE, PE-CF594, PE-Cy7, eFluor 660, APC-R700, APC-Cy7 |
| OMIP-058 | 355, 405, 488, 532, 628 | FITC, Brilliant Blue 630, Brilliant Blue 660, Brilliant Blue 700, Brilliant Blue 790, PE, PE-CF594, PE-Cy5, PE-Cy5.5, PE-Cy7, APC, Alexa Fluor 700, APC-Cy7, BUV395, BUV496, BUV563, BUV661, BUV737, BUV805, LIVE-DEAD Fixable Blue, Brilliant Violet 421, Brilliant Violet 510, Brilliant Violet 570, Brilliant Violet 605, Brilliant Violet 650, Brilliant Violet 711, Brilliant Violet 750, Brilliant Violet 786 |
| OMIP-059 | 355, 405, 488, 561, 640 | BUV395, Brilliant Violet 421, APC, APC-Cy7, PE, Brilliant Blue 515, PerCP-Cy5.5 |
| OMIP-060 | 355, 405, 488, 532, 628 | FITC, Brilliant Blue 630, Brilliant Blue 660, Brilliant Blue 700, Brilliant Blue 790, PE, Alexa Fluor 594, PE-Cy5, PE-Cy5.5, PE-Cy7, Alexa Fluor 647, Alexa Fluor 700, APC-H7, BUV395, BUV496, BUV563, BUV661, BUV737, BUV805, LIVE-DEAD Fixable Blue, Brilliant Violet 421, Brilliant Violet 480, Brilliant Violet 570, Brilliant Violet 605, Brilliant Violet 650, Brilliant Violet 711, Brilliant Violet 750, Brilliant Violet 786 |

| | | |
|---|---|---|
| OMIP-061 | 355, 405, 488, 532, 628 | PerCP-eFluor 710, Brilliant Violet 421, Brilliant Violet 510, Brilliant Violet 570, Brilliant Violet 605, Brilliant Violet 650, Brilliant Violet 711, Brilliant Violet 750, Brilliant Violet 786, PE, PE-CF594, PE-Cy5, PE-Cy7, BUV395, BUV496, BUV661, BUV737, APC, APC-R700, LIVE-DEAD Fixable Blue |
| OMIP-062 | 405, 488, 561, 640 | Brilliant Violet 421, Brilliant Violet 510, Brilliant Violet 605, Brilliant Violet 650, Brilliant Violet 711, Brilliant Violet 785, FITC, PerCP-Cy5.5, PE, PE-Dazzle 594, PE-Cy7, Alexa Fluor 647, Alexa Fluor 700, LIVE-DEAD Fixable Near-IR |
| OMIP-063 | 355, 402, 442, 488, 561, 628 | Alexa Fluor 488, Brilliant Blue 630, Brilliant Blue 660, Brilliant Blue 700, Brilliant Blue 790, PE-CF594, PE-Cy5, PE-Cy7, APC, APC-R700, APC-Cy7, BUV395, BUV496, BUV563, BUV615, BUV661, BUV737, BUV805, LIVE-DEAD Fixable Blue, Brilliant Violet 421, Brilliant Violet 480, Brilliant Violet 570, Brilliant Violet 605, Brilliant Violet 650, Brilliant Violet 711, Brilliant Violet 750, Brilliant Violet 786, BYG584 |
| OMIP-064 | 637, 532, 488, 405, 355 | BUV395, BUV496, BUV563, BUV661, BUV737, BUV805, Brilliant Violet 421, Brilliant Violet 480, Brilliant Violet 570, Brilliant Violet 605, Brilliant Violet 650, Brilliant Violet 750, Brilliant Violet 786, Brilliant Blue 660, LIVE-DEAD Fixable Blue, FITC, PerCP-Cy5.5, Brilliant Violet 711, Brilliant Blue 790, PE, PE-CF594, PE-Cy5, PE-Cy5.5, PE-Cy7, Alexa Fluor 647, Alexa Fluor 700, APC-Fire 750 |
| OMIP-065 | 405, 488, 561, 640 | eFluor 450, LIVE-DEAD Fixable Aqua, APC, Brilliant Violet 786, PE, FITC, APC-eFluor 780, PE-Cy7, PerCP-eFluor 710, PE-CF594, Alexa Fluor 700, SuperBright 600, SuperBright 645, SuperBright 702 |
| OMIP-066 | 488, 405, 532, 355, 628 | VioBright 515, Brilliant Blue 700, Brilliant Violet 421, Brilliant Violet 510, SuperBright 702, Brilliant Violet 786, PE, PE-Dazzle 594, PE-Cy7, BUV395, BUV805, Alexa Fluor 647, APC-R700, APC-Fire 750, LIVE-DEAD Fixable Blue |
| OMIP-067 | 355, 405, 488, 532, 628 | Alexa Fluor 700, BUV496, BUV805, PE, Brilliant Blue 700, Brilliant Violet 786, Brilliant Violet 711, BUV661, Brilliant Violet 570, Brilliant Blue 790, PE-Cy7, Brilliant Violet 605, Brilliant Violet 650, PE-Dazzle 594, BUV563, PE-Cy5.5, BUV395, BUV737, Brilliant Violet 421, Brilliant Blue 630, Alexa Fluor 647, Brilliant Blue 660, APC-Vio770, PE-Cy5, FITC, VioGreen, Brilliant Violet 750, LIVE-DEAD Fixable Blue |

| | | |
|---|---|---|
| OMIP-069 | 355, 405, 488, 561, 640 | BUV395, BUV496, BUV563, BUV615, BUV661, BUV737, BUV805, LIVE-DEAD Fixable Blue, Brilliant Violet 421, Brilliant Violet 480, Brilliant Violet 510, Brilliant Violet 570, Brilliant Violet 605, Brilliant Violet 650, Brilliant Violet 711, Brilliant Violet 750, Brilliant Violet 785, SuperBright 436, eFluor 450, Pacific Orange, Brilliant Blue 515, FITC, PerCP, PerCP-Cy5.5, PerCP-eFluor 710, Spark Blue 550, PE, PE-Dazzle 594, PE- Alexa Fluor 610, PE-Cy5, PE-Alexa Fluor 700, PE-Cy7, PE-Fire 810, APC, Alexa Fluor 647, Spark NIR 685, APC-R700, APC-H7, APC-Fire 810 |
| OMIP-070 | 355, 405, 488, 532, 628 | LIVE-DEAD Fixable Blue, BUV395, BUV496, BUV563, BUV661, BUV737, BUV805, Brilliant Violet 421, Brilliant Violet 480, Brilliant Violet 570, Brilliant Violet 605, Brilliant Violet 650, Brilliant Violet 711, Brilliant Violet 750, Brilliant Violet 786, FITC, Brilliant Blue 630, Brilliant Blue 660, Brilliant Blue 700, Brilliant Blue 790, PE, PE-CF594, PE-Cy5, PE-Cy5.5, PE-Cy7, eFluor 660, Alexa Fluor 700, APC-H7 |
